# The clock transcription factor BMAL1 is a key regulator of extracellular matrix homeostasis and cell fate in the intervertebral disc

**DOI:** 10.1101/2023.02.12.528214

**Authors:** Michal Dudek, Honor Morris, Natalie Rogers, Dharshika RJ Pathiranage, Danny Chan, Karl E Kadler, Judith Hoyland, Qing-Jun Meng

## Abstract

The circadian clock in mammals temporally coordinates physiological and behavioural processes to anticipate daily rhythmic changes in their environment. Chronic disruption to circadian rhythms (e.g., through ageing or shift work) is thought to contribute to a multitude of diseases, including degeneration of the musculoskeletal system. The intervertebral disc (IVD) in the spine contains circadian clocks which control ∼6% of the transcriptome in a rhythmic manner, including key genes involved in extracellular matrix (ECM) homeostasis. However, it remains largely unknown to what extent the local IVD molecular clock is required to drive rhythmic gene transcription and IVD physiology. In this work, we identified profound age-related changes of ECM microarchitecture and an endochondral ossification-like phenotype in the annulus fibrosus (AF) region of the IVD in the *Col2a1*-*Bmal1* knockout mice. Circadian time series RNA-Seq of the whole IVD in *Bmal1* knockout revealed loss of circadian patterns in gene expression, with an unexpected emergence of 12-hour ultradian rhythms, including FOXO transcription factors. Further RNA sequencing of the AF tissue identified region-specific changes in gene expression, evidencing a loss of AF phenotype markers and a dysregulation of ECM and FOXO pathways in *Bmal1* knockout mice. Consistent with an up-regulation of FOXO1 mRNA and protein levels in *Bmal1* knockout IVDs, inhibition of FOXO1 in AF cells suppressed their osteogenic differentiation. Collectively, these data highlight the importance of the local molecular clock mechanism in the maintenance of the cell fate and ECM homeostasis of the IVD. Further studies may identify potential new molecular targets for alleviating IVD degeneration.

## Introduction

Evolutionarily conserved circadian (∼24-hourly) rhythms persist throughout biology, with almost every aspect of our physiology and behaviour having evolved around the rotation of the earth. Endogenous circadian rhythms define our daily patterns of sleep, eating and exercise, moderate the scale of an immune response we mount, determine how our body responds to medications, and gate daily patterns of metabolism^1^. Evolved as a homeostatic mechanism, this temporal alignment of behaviour and physiology to the external environment ensures optimisation of metabolism and energy allocation to anticipate daily variations in physiological demands^2^. In a tissue context-dependent manner, circadian clocks drive 24-hour rhythmicity in approximately 5–10% of the genome,^3^ 5–20% of the proteome^4–6^ and ∼25% of the phosphoproteome^7^. Like many other regulatory processes, the circadian clock changes during ageing, losing its precise temporal control with the robustness of circadian rhythms, in terms of oscillatory amplitude and circadian phase, declining with age in both animal models and humans^8^. As such, the functional decline of circadian rhythms has been proposed as a potential mechanism driving an increased risk of various diseases, including metabolic syndromes, cancer and musculoskeletal conditions^8–10^. Indeed, genetic models of circadian disruption exhibit age-related musculoskeletal conditions including osteoporosis, osteoarthritis and tendinopathy^11–14^. In humans, prolonged misalignment of internal circadian rhythms with environmental rhythms, such as those seen in chronic shift workers or frequent long-haul travellers, is associated with profound consequences for health and well-being^15,16^. Given that both population ageing and chronic circadian misalignment are increasingly prevalent, understanding the importance of biological timing mechanisms in age-related diseases is becoming more and more relevant.

Intervertebral discs (IVDs) are extracellular matrix (ECM)-rich fibrocartilaginous joints that allow flexion and shock absorption in the spine. They consist of an outer fibrous ring, the annulus fibrosus (AF), which surrounds an inner gelatinous centre, the nucleus pulposus (NP) rich in proteoglycans. The AF consists of several weaved layers (laminae) of tendon-like tissue composed predominantly of collagen. Correct arrangement of the layers as well as the collagen fibres is essential to counteract mechanical forces and contain the NP when the spine is subject to mechanical load. The AF consists of distinct regions — the inner AF, adjacent to the NP and rich in type II collagen, and an outer AF to the peripheral edge of the disc consisting primarily of type I collagen. The inner AF is populated by rounded fibrocartilage cells, while cells in the outer AF show a more fusiform fibroblast-like morphology. These cells are organised in a linear arrangement, with the degree of linear alignment increasing as proximity to the NP decreases. With age, there is progressive degeneration of the IVD, which is reported to affect approximately 90% of individuals over the age of 50 years^17^. This degeneration is characterised by reduced tissue hydration (due to loss of proteoglycan), fibrosis, calcification and altered cell population and morphology^18–23^. These pathological changes form a vicious circle whereby alterations in cell behaviour and matrix composition impair tissue mechanics (and ultimately tissue function), which in turn drive an exaggerated and aberrant cellular response.

Despite the notion of daily variations in spine physiology being recorded as early as 1724, evidence of an intrinsic circadian rhythm within cells of the IVD was only provided recently ^24^. The molecular circadian clock within the mouse IVD drives rhythmic IVD transcription and shows strong tissue-specificity, with little overlap even within the musculoskeletal systems. These rhythmic outputs include genes relevant to ECM turnover (e.g., *Adamts1, Timp4, Serpinh1 and Itgb1*) and endoplasmic reticulum (ER) stress (e.g., *Pak1, Atf6 and Chac1*) and are believed to be essential for coping with the daily stresses that the IVD is exposed to within its unique physiological niche^24^. Degeneration of the IVD is a complex and multifactorial process associated with ageing. Significantly, these IVD clocks are disrupted by ageing and inflammation, two known risk factors for IVD degeneration, suggesting the involvement of clock mechanisms in driving a predisposition to the development of acclerated ageing and degeneration in the IVD^24^. However, it remains largely unknown how the IVD clock regulates ECM homeostasis and cell fate in the IVD, and what the underlying molecular links are.

Here, a circadian clock disruption model (*Col2a1*-*Bmal1* knockout mouse) was analysed using atomic force microscopy and electron microscopy, and time-resolved and tissue-type specific transcriptomic analysis, to investigate the importance of circadian clocks in the homeostasis and degeneration of the ECM-rich intervertebral disc tissues. Our data reveal a profound age-related degeneration characterised by dysregulated ECM microarchitecture in the *Bmal1* KO IVDs, with a major osteogenic component linked to the emergence of a putative ultradian 12-hour clock. The circadian transcriptome of the whole disc as well as a comparison of the AF transcriptome indicate an apparent loss of AF phenotype markers and altered ECM homeostasis, with FOXO genes identified as key drivers of the *Bmal1* KO phenotype.

## Results

### Tissue-specific knockout of Bmal1 predisposes the IVD to age-related extracellular matrix degeneration

Masson’s trichrome staining of the *Col2a1*-*Bmal1* knockout (KO) mouse showed age-related intervertebral disc phenotypes, characterised by thinning of the cartilage end plate and bony protrusions into the growth plate (Fig. S1), consistent with our earlier report using standard histology and X-ray imaging^25^. To gain further insight into changes to the ECM architecture and cell morphology, a newly developed RGB-trichrome staining method^26^ was employed (Figure 1A). This histology analysis allows for clear differentiation not only between tissue types, but also between calcified and uncalcified ECM. Wildtype (WT) IVDs showed clear demarcation between IVD tissue components, with the collagen-rich AF staining in the red spectrum and more proteoglycan-rich tissues such as the NP staining blue. While the lamellae of the inner AF were still evident in the *Bmal1* KO tissue, the resident AF cells appear rounder in morphology when compared with the flattened fibroblast-like cells typical of the healthy AF. The lamellar structure of the outer AF was progressively lost, and large rounded cells appeared. These cells appeared to have migrated outwards towards the periphery of the outgrowth, evidenced by the brush stroke-like pattern of dark blue staining that radiated outwards from the lamellar AF (Figures 1A and B). This blue region then transitions into a distinct yellow-green matrix (Figure 1B(4)), likely representing calcified ECM^26^. This cartilaginous structure is visible macroscopically upon dissection of the discs (Fig. S2). It appeared that this tissue was derived from rounded hypertrophic chondrocyte-like cells in the KO AF (Figure 1B). This change of cellular phenotype was reiterated by scanning electron microscopy, showing large, rounded cells with electron dense cytoplasmic compartments in the outer AF of *Bmal1* KO (Figure 1C). Segmentation of the serial electron microscopy images revealed striking difference in cell morphology. The *Bmal1* KO cells appeared polarised with numerous protrusions on one side of the cell and rounded plasma membrane on the other. Some of the protrusions containing electron dense material became disconnected from the main body as if left behind by a migrating cell (Figure 1D, Figure S3 and Supplementary videos 1 and 2).

**Figure 1.**
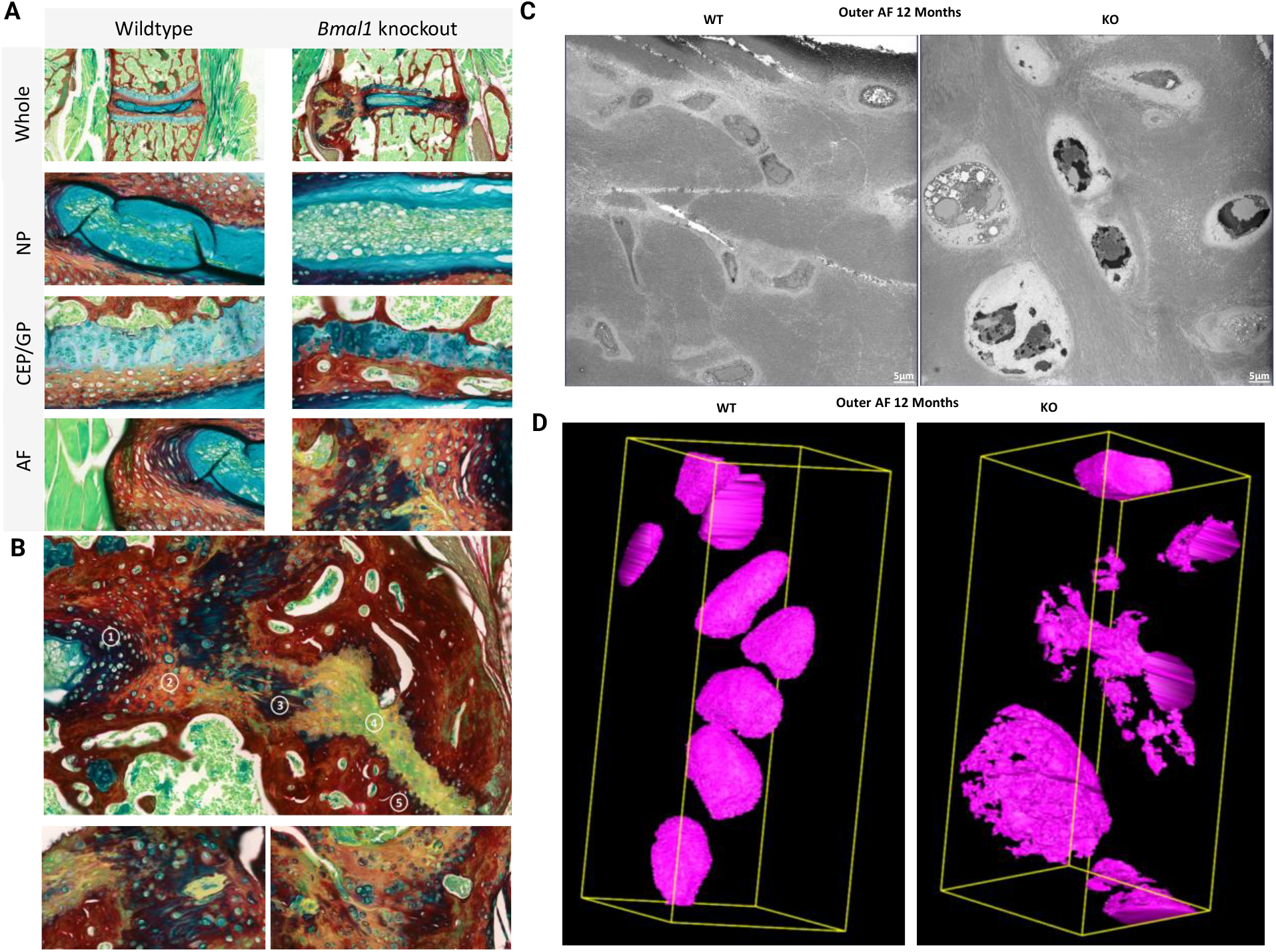
Col2a1-specific *Bmal1* KO leads to age related IVD degeneration. **A**. RGB trichrome staining of 12-month-old wildtype versus Col2a1-*Bmal1* KO lumbar intervertebral disc. **B**. Representative heterotopic ossification region of the aged Col2a1*-Bmal1* KO IVD. Top, Composite image of 10x magnification snapshots, stitched together to visualise the whole region of heterotopic ossification. Numbers represent a proposed migratory route of cells originating from the inner AF (1), moving to the outer AF (2), the proteoglycan-rich region fanning out from the IVD (3), the non-calcified cartilage region (4) and finally cartilage-bone interface showing pits with resident hypertrophic chondrocyte-like cells (5). Bottom left, x40 magnification showing in detail additional representative staining patterns similar to (3) in the top panel. Staining appears to show a brushstroke-like effect from the centre of the disc (right to the peripheral edge (left, yellow-staining region). Bottom right, x40 magnification showing in greater detail a representative region similar to (4, 5) in the top panel. Cartilage-bone interface is visible, with blue-staining rounded cells clustered around the interface. **C**. Representative images from scanning electron microscopy (SEM) showing WT and *Bmal1* KO outer AF tissues at 12 months of age. **D**. 3D rendering of segmented cells from serial block-face SEM from WT and *Bmal1* KO outer AF tissues at 12 months of age.

### Dysregulated collagen organisation and nano-architecture in the IVD of Bmal1 KO mice

Given the disruption to the lamellar arrangement of the collagen-rich AF in *Bmal1* KO mice, AFM and EM were used to examine collagen architecture on the nanoscale. The inner AF of young mice (6 weeks old) showed similar collagen arrangements and typical D period length of 67 nm between WT and *Bmal1* KO (Figure 2A). However, fibrils showed a significantly thinner mean diameter of 85.12(±19.73) nm in *Bmal1* KO tissue as compared with 90.45(±23.28) nm in wildtype (P<0.001, Figure 2B). Moreover, there was a negative shift in the distribution of fibril diameters, with no fibrils of ≥150nm identified in *Bmal1* KO tissue (Figure 2B). Collagen fibrils in the ageing WT inner AF were arranged in a linear fashion, as visualised by AFM at coronal and EM at transverse planes (Figure 2C). Fibrils in the *Bmal1* KO inner AF appeared slightly disorganised both by AFM and EM imaging (Figure 2C). Thinning of collagen fibrils was evident in *Bmal1* KO animals, with these fibrils having a significantly smaller mean diameter of 101(±20.62) nm compared with 156.8(±24.2) nm in wildtype (P<0.0001, Figure 2D). Plotting the distribution of fibril diameters showed a leftwards shift in diameter in *Bmal1* KO fibrils. Length of the D period showed no significant change (Figure 2D). The outer AF of ageing *Bmal1* KO animals showed a more profound change than the inner AF. Linear fibril arrangement was maintained in the outer AF of the wildtype disc. However, visible collagen architecture was almost unidentifiable in *Bmal1* KO (Figure 2E). The tissue was also pitted by numerous small indentations and mounds, which may have been left by calcification deposits (Figure 2E).

**Figure 2.**
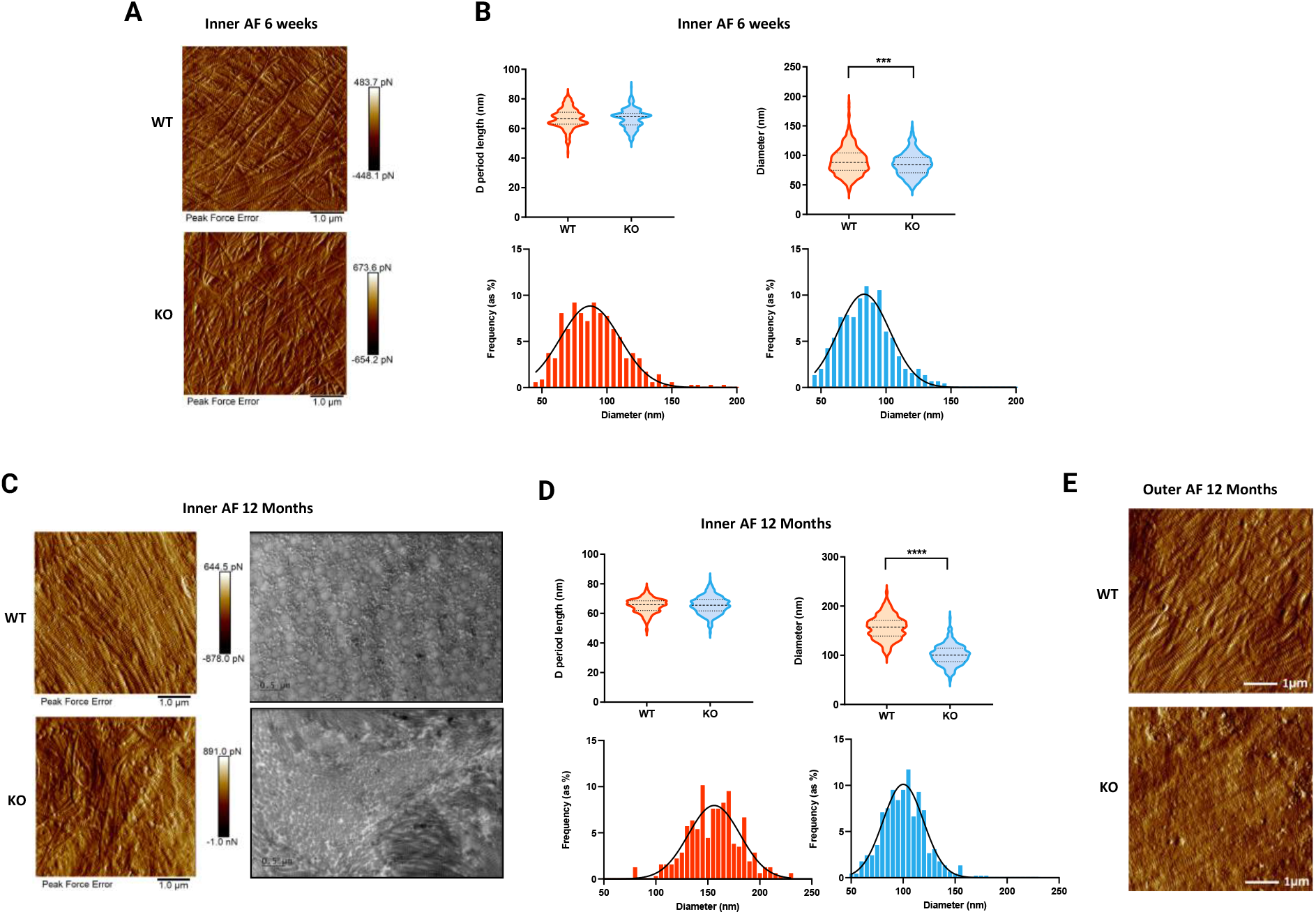
AFM analysis of the AF in mouse lumbar intervertebral discs reveals thinner and disorganised collagen fibres in the Col2a1-*Bmal1* KO. **A**. Representative peak force error AFM images of WT and *Bmal1* KO inner AF at 6 weeks of age showing collagen fibril morphology. **B**. Quantification of the length of fibril D periods and diameter from AFM images of WT and *Bmal1* KO inner AF at 6 weeks of age. **C**. Representative peak force error AFM images of WT and *Bmal1* KO inner AF at 12 months of age showing morphology of collagen fibrils. **D**. Quantification of the length of fibril D periods and diameter from AFM images of wildtype and *Bmal1* KO inner AF at 12 months of age. **E**. Representative peak force error AFM images of wildtype and *Bmal1* KO outer AF at 12 months of age showing morphology of collagen fibrils. Tissues from three animals per genotype and per condition were imaged and quantified. Student’s t test, *** - p <0.001.

### Outer AF of the Bmal1 KO IVD undergoes endochondral ossification

Given the apparent deposition of cartilage and bone-like ECM within the outgrowth region of the *Bmal1* KO IVDs, resident cells were examined to gain further insight into their phenotype. Their rounded morphology and cartilaginous territorial matrix led to the hypothesis that these cells resemble hypertrophic chondrocytes. This hypothesis was supported by immunohistochemistry (IHC) staining for chondrogenic makers such as ALPL and SOX9 (Figure 3). ALPL-positive cells were generally absent within the WT AF, however, were abundant in *Bmal1* KO animals. These cells were present mostly within the central cartilaginous region and peripheral edges of the outgrowth and showed a rounded morphology (Figure 3, Fig. S4). SOX9 was expressed in the wildtype AF, but SOX9-positive cells appeared to be more abundant in the *Bmal1* knockout IVD, particularly on the peripheral edge of the outgrowth (Figure 3, Fig. S4).

**Figure 3.**
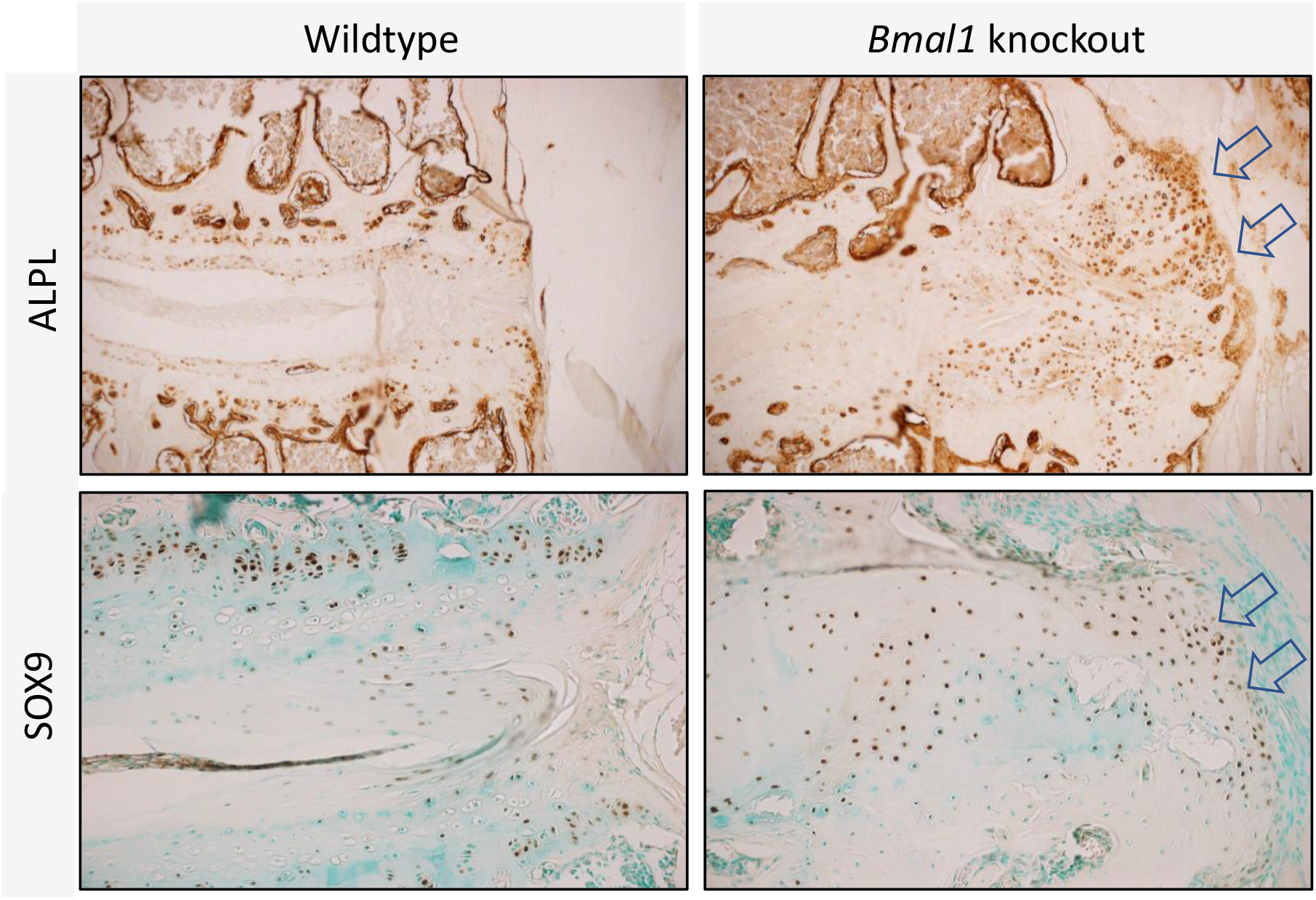
Aged *Bmal1* KO AF expresses increased levels of hypertrophic chondrocyte markers. Representative immunohistochemistry staining of hypertrophic chondrocyte markers ALPL and SOX9 in 12-month-old WT and Col2a1-*Bmal1* KO mice. Arrows highlight expression of ALPL and SOX9 in the peripheral edge of the heterotopic ossification region in 12-month-old Col2a1-*Bmal1* KO lumbar intervertebral discs. ALPL and SOX9-positive cells are prolific throughout this region, particularly towards the peripheral edges. N=3.

### Comparative analysis of the WT and Bmal1 KO circadian transcriptome revealed emergence of 12-hour rhythmic genes in KO

*Bmal1* is an essential component of the circadian clock mechanism without which the 24-hour oscillations patterns in gene expression will be abolished. To fully assess the effect of IVD-specific *Bmal1* knockout, we performed circadian time series RNAseq in *Bmal1* KO whole IVDs and performed a comparative analysis with our published WT mouse IVD data^25^. This analysis identified 1092 genes (approximately 6% of the IVD transcriptome) that cycled with a circadian pattern (Figure 4A) in WT. GO term analysis of circadian genes showed over-represented terms including ‘fatty acid metabolic process’, ‘circadian regulation of gene expression’, ‘response to endoplasmic reticulum stress’ and ‘proteasomal protein catabolic process’ (Figure 4B). It is of particular interest to note the circadian expression of genes relating to ECM homeostasis and chondrocyte/AF cell physiology, such as *Sost, Ihh, Tgfβi*, collagen family genes and *Adamts* family genes (Supplementary Table 1). Unsurprisingly, the circadian expression patterns of these transcripts were lost in the *Bmal1* knockout IVDs, supporting the role of BMAL1 as a critical regulator of IVD circadian rhythmicity.

**Figure 4.**
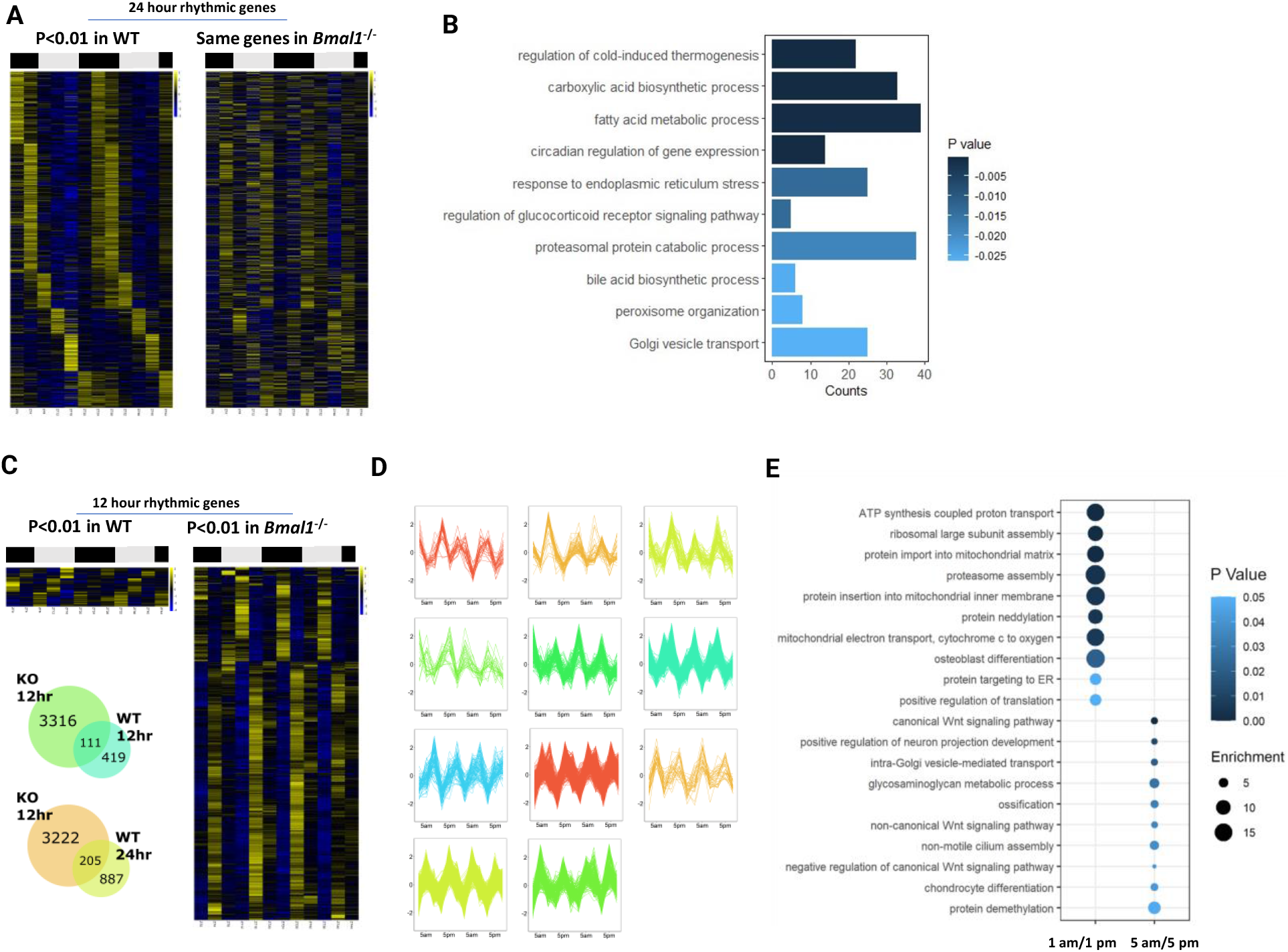
Emergence of 12-hour ultradian genes in *Bmal1* KO IVDs. **A**. Heatmaps depicting the expression patterns of the 1,092 rhythmic genes (P<0.01) in WT (left). Equivalent expression patterns of these genes were shown for the *Col2a1*-*Bmal1* KO IVD (right). Genes were organised according to timing of peak expression. White bars represent the day; black bars represent the night. **B**. GO term enrichment analysis of the circadian transcriptome, showing the top 10 terms according to P value. **C**. Heatmaps depicting the expression patterns of the ultradian transcriptome in WT and *Bmal1* KO IVDs and Venn diagram showing overlap between *Bmal1* KO and WT ultradian transcriptome, and *Bmal1* KO ultradian and WT circadian transcriptome. **D**. Waveform clustering of ultradian transcripts expressed in the *Bmal1* KO mouse intervertebral disc. Y axis values show normalised expression. **E**. GO enrichment analysis of the ultradian transcriptome in the *Bmal1* KO IVDs, segregated by the time of oscillation peak. The top 10 enriched terms were displayed, ranked by P value.

Somewhat surprisingly, a profound ultradian rhythmic pattern in transcriptional dynamics was identified in the *Bmal1* knockout (Figure 4C). 3,427 genes exhibited a 12-hour oscillatory period (p<0.01). This represents a substantial 17% of the total transcripts identified in the KO mouse IVD. 12-hour gene expression patterns were less apparent within the WT disc (530 genes at p<0.01). Among these, only 111 genes were common to both genotypes. 3,316 genes therefore showed *de novo* ultradian oscillations in *Bmal1* knockout (Figure 4C). The majority of these ultradian genes did not show 24-hour rhythms in WT IVDs (Fig. 4C). This implies a profound switch in temporal gene expression dynamics upon disruption of the molecular circadian clock.

Clustering of the ultradian transcriptome by temporal pattern produced 11 clusters with robust oscillatory patterns (Figure 4D). Of note, these ultradian genes can be largely segregated into two ‘time of peak’ groups. Approximately 70% of the ultradian transcriptome showed expression peaks at 5 am and 5 pm, while the remaining 30% of genes showed peaks in their expression at 1am and 1pm. GO term analysis revealed a clear distinction between the two groupings (Figure 4E). 5 am/5 pm peaking genes showed significant enrichment for processes relating to matrix and cell-cell signalling such as ‘glycosaminoglycan metabolic process’, ‘non-canonical Wnt signalling pathway’, ‘ossification’, ‘cartilage development’, ‘osteoblast differentiation’ and ‘regulation of cell response to growth factor stimulus’, with a list of collagen family genes (Figure 4E; Fig. S5, Fig. S6). In contrast, 1 am/1 pm peaking genes showed a strong enrichment for terms pertaining to the electron transport chain subunits, mitochondrial translation, cytoplasmic translation, protein transportation and proteasome assembly (Figure 4E). Of the 88 nuclear-derived genes encoding Complexes I–X of the electron transport chain, a striking 72% were found to be significantly ultradian. None of these genes show a significant ultradian pattern in the WT disc. Electron microscopy imaging of mitochondria from WT and *Bmal1* KO AF showed electron dense inclusion bodies in KO tissues, suggesting that gene expression changes may indeed have effects on the structure or functioning of this organelle (Fig. S7).

### Promoter analysis of ultradian genes and AF-specific transcriptomics implicate FOXO transcription factors as effectors for the osteogenic AF phenotype

UPR analysis of genes peaking at 5 am/5 pm showed strong enrichment of forkhead box (FOX) factor motifs (FOXA1, FOXD3, FOXI1, FOXO3, FOXQ1), ARID3A, CEBPA, HOXA5, NFATC2, NOBOX and SRY (Figure 5A). 11 out of 25 Fox genes expressed in the mouse IVD showed significant ultradian rhythmicity in the *Bmal1* knockout disc (Figure 5B, Fig.S8). Genes peaking at 1 am/1 pm were enriched for motifs annotated to ELK4, IRF2, NR1H2:RXRA, NR3C1, SRF and ZNF143 (Fig. S9).

**Figure 5.**
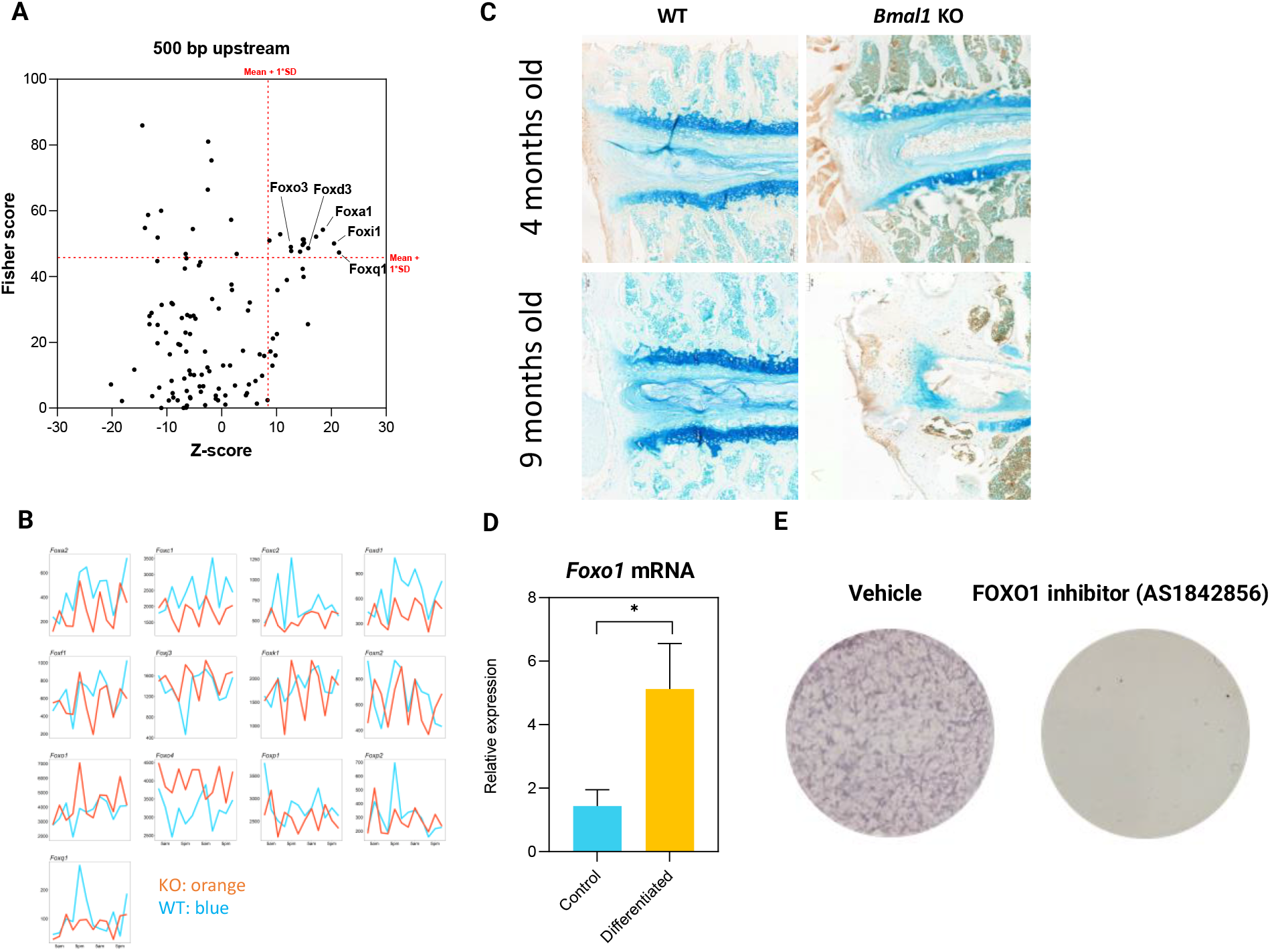
FOXO1 is involved in the development of the degenerative IVD phenotype in *Bmal1* KO mice. **A**. Scatter plot of individual transcription factor binding sites (TFBPS) plotted against the Fisher scores and Z-score rankings of their enrichment in the regions 500 bp upstream of the transcription start sites of 5 am/5 pm peaking genes in the ultradian transcriptome. **B**. Traces of WT (blue) and *Bmal1* KO (orange) gene expression of FOX transcription factor genes showing significant ultradian rhythmicity in the *Bmal1* KO IVDs. **C**. Immunohistochemistry staining of FOXO1 in 4- and 9-month-old wildtype and Col2a1-*Bmal1* KO mice. Arrows highlight positive expression of FOXO1 in the heterotopic ossification region. **D**. *Foxo1* gene expression in AF-S cells following 3 weeks of culture in osteogenic or control medium (N=3). **E**. Alkaline phosphatase staining of AF-S cells following 3 weeks of culture in osteogenic medium in the presence or absence of 0.5 µM a FOXO1 inhibitor AS1842856 (N=3).

To gain more insight into the AF phenotype, we performed comparative RNAseq on AF tissues dissected at 8 am from WT and *Bmal1* KO mice. 365 genes were differentially expressed (63% downregulated in *Bmal1* knockout and 37% upregulated) between WT and *Bmal1* KO AF (Fig. S10A). Enrichment analysis identified pathways related to ECM organisation, tissue development and FOXO target genes (Fig. S10B). Differentially expressed genes included key regulators of IVD and cartilage homeostasis, such as *Ccn2* (also known as *Ctgf*), *Cnmd, Foxa2, Gdf6, Grem1, Gli1* and *Ptch1*. Interestingly, many genes described as putative AF marker genes were downregulated in *Bmal1* KO AF tissue, including *Fbln1, Fbln2, Col1a1, Col1a2, Chad* and *Comp*. This is indicative of a loss of AF phenotype in *Bmal1* knockout and supports a transition of AF cells to a more hypertrophic chondrocyte-like phenotype in *Bmal1* knockout.

Among the FOXO factors, FOXO1 has been reported to play a role in age-related IVD degeneration^27,28^, making it a key candidate for further investigation. *FOXO1* showed a strong 12-hour rhythmic expression pattern in *Bmal1* KO IVDs (P<0.001), with much higher expression levels at the 5 pm peaks (Figure 5B). As such, protein levels of FOXO1 were assessed using IHC. In 4 month- and 9 month-old WT mouse IVDs, FOXO1 signals were almost undetectable. In contrast, in age-matched *Bmal1* KO IVDs, FOXO1 expression was readily detected in the AF region, particularly in the outer AF where osteogenic differentiation occurs (Figure 5C). In an *in vitro* osteogenic differentiation model of human AF cells, *FOXO1* mRNA levels showed an approximately 3-fold induction following osteogenic differentiation, implying *FOXO1’s* involvement in osteogenic differentiation of AF cells (Figure 5D). Consistent with this notion, pharmacological inhibition of FOXO1 with AS1842856 led to an inhibition of osteogenic differentiation in human AF cells (Figure 5E). Taken together, these data support a significant role of FOXO1 in mediating the osteogenic AF phenotype in *Bmal1* KO IVDs.

## Discussion

In this study, using *Col2a1*-specific knockout of the essential circadian clock component *Bmal1*, we investigated the effect of circadian clock disruption on the tissue architecture and gene expression profiles of the intervertebral discs. Profound age-related IVD degeneration was identified in the AF region of the knockout animals; further analysis using electron microscopy and atomic force microscopy identified changes in cell phenotype as well as disorganisation of extracellular matrix structure. Analysis of the IVD circadian transcriptome revealed a loss of the 24-hour rhythmic gene patterns and emergence of significant ultradian 12-hour oscillation patterns in the *Bmal1* knockout IVDs. Both the circadian transcriptome of the whole disc as well as a comparison of the AF transcriptome pointed to ECM homeostasis as a major *Bmal1* target pathway, revealing FOX genes as key drivers of the *Bmal1* KO phenotype. Importantly, our data demonstrate FOXO1 as a critical *Bmal1*-regulated transcription factor involved in the osteogenic differentiation of human AF cells.

Despite the highly conserved expression of core clock genes, cross tissue comparisons of circadian functions have revealed an amazing degree of tissue specificity, both in terms of the input pathways to the clock, and output clock-controlled targets^3,29^. As shown previously and in this work, the molecular circadian clock within the WT mice drives rhythmic transcription of ∼6% of the IVD transcriptome and shows strong tissue-specificity. These rhythmic outputs are thought to be essential for coping with the daily stresses that the IVD is exposed to within its unique physiological niche^30–32^. The majority of identified circadian genes cycled with a peak expression during waking hours. This ‘active phase’ group consisted of genes relevant to ECM turnover (e.g., *Adamts1, Timp4, Serpinh1, Itgβ1*) and ER stress (e.g., *Pak1, Atf6, Chac1*), among others.

Age-related degeneration of the intervertebral disc is a multi-factorial process, involving a complex interplay between discrete tissue subtypes within the IVD. We have previously shown that ageing disrupts circadian rhythms in IVDs and that the *Bmal1* KO IVD phenotype resembles some of the features seen in aged mice and IVD degeneration^25^. Disruption to the collagen architecture of the ECM is a hallmark of age-related degeneration in cartilaginous tissues. Collagen permits structural flexibility to the disc, necessary for coping with compressive, tensile and shear stresses. In the non-degenerative disc, the AF consists mainly of thick type I collagen fibres with a well-defined linear arrangement. With ageing, the collagen content of the AF declines^33^, thinner fibres become more prevalent^34^ and linear arrangement is disrupted^35^. These changes are consistent with a shift to a more cartilage-like collagen structure^36^. Our finding that collagen fibrils within the inner AF are thinner in *Bmal1* KO animals as early as 6 weeks (before the onset of observable disc degeneration) suggests that the inner AF could be the primer for later degeneration. We have previously reported that disruption of the circadian clock in tendons leads to malfunctioning of the secretory pathway, and causes abnormal collagen fibrils and collagen accumulation^37^. As a tissue of comparable composition and structure, a similar mechanism could be at play in the AF of intervertebral discs. Thinner collagen fibrils may render the AF less resilient to mechanical strain, leading to accelerated damage or an exaggerated repair response.

An exaggerated cell repair response could explain the cartilaginous protrusion and cell morphology changes observed in the outer AF. Although we have not been able to determine the origin of these cells, studies using single cell RNA-sequencing have identified multiple cell type subsets within the healthy NP and AF, including progenitor-like cells in both tissues^38,39^. Moreover, AF cells have been shown to be capable of differentiating into chondrocytes and osteoblasts *in vitro* as well as *in vivo, driving ossification in rat lumbar disc needle puncture model*^23^. Migration of progenitor-like cells within the IVD tissue has also been identified previously, such as the migration of cells from a growth plate-adjacent niche through the AF into the centre of the disc^40,41^. Therefore, in the AF of *Bmal1* KO, transition of local cells or invasion by neighbouring regions are equally plausible.

In line with the AFM and EM findings of changes in collagen matrix composition, the circadian transcriptome of WT IVDs showed ‘response to endoplasmic reticulum stress’ and ‘Golgi vesicle transport’ as among the overrepresented circadian regulated processes. Rhythmicity in most of the genes belonging to these categories was lost in the *Bmal1* KO discs. Surprisingly, a new ultradian pattern of expression emerged in the KO IVDs with two antiphase 12-hour oscillations. Among the 5 am/5 pm peaking genes we revealed overrepresented processes such as ‘intra-Golgi vesicle mediated transport’, ‘glycosaminoglycan metabolic process’ and ‘ossification’, suggesting dysregulation of protein maturation and secretion processes. In WT liver tissues, ultradian rhythms have been referred to as ‘harmonics’ of the circadian rhythm, i.e., a component frequency of the 24-hour circadian oscillation. This implies that ultradian rhythms and circadian rhythms could share a common origin^42^. However, this mechanism is unlikely to be at play in the KO datasets presented here due to the molecular defects of the 24-hour circadian clocks. Genes encoding components of the circadian clock for the most part did not show a shift towards a 12-hour pattern in *Bmal1* knockout IVDs, with the exception of *Npas2*, a binding partner of BMAL1 to activate E-box containing genes (Figure S11). Indeed, promoter analysis for over-represented transcription factor binding sites did not identify the E-box motif to be significantly enriched within the ultradian dataset. This suggests that alternative oscillators may be at play in the *Bmal1* knockout IVD to drive ultradian gene transcription.

The FOX family, consisting of 44 transcription factors in humans and mice, regulate diverse cellular functions, from mitochondrial homeostasis and stress response to cell fate and differentiation^43^, through binding of the highly conserved forkhead box domain. While few studies have directly implicated a role for FOX signalling in osteogenic differentiation^44^, the role of FOX signalling in ageing, longevity and regeneration processes bears relevance to these studies. The FOXO subfamily is of particular interest due to its role in lifespan determination and the pathogenesis of age-related diseases^45,46^. Recently, FOXO genes have been implicated in age-related IVD degeneration^27,28^. Though FOX factors have not yet been explicitly implicated in ultradian rhythmicity, FOXO1 was identified as a predicted upstream regulator of circatidal rhythms in the intertidal mollusc *C. rota*^47^. Given the commonality between these findings, the conservation of ultradian rhythms previously described, enrichment of FOX binding motifs in the ultradian transcriptome as well as ultradian rhythmicity of many FOX family genes, FOX transcription factors may represent a putative regulator of ultradian transcriptional rhythms in the *Bmal1* KO IVDs.

Collectively, our findings implicate circadian rhythms and the core clock transcription factor *Bmal1* as critical regulators of intervertebral disc physiology. Given the increasing expansion of the ageing populations in a modern world which antagonises our innate circadian timing mechanisms, the circadian regulation of ECM homeostasis and IVD cell fate identified here may provide a new molecular mechanism and potential therapeutic targets for age-related IVD degeneration and low back pain.

## Acknowledgments

We thank J Takahashi (UT Southwestern Medical Center, US) for the PER2::Luc mouse line. We thank GC Van Den Akker, TJ Welting and JW Voncken (Maastricht University, Netherlands) for their kind provision of the human AF-S cell line. We thank the Genomics Core Facility (A Hayes and L Zeef) for their kind assistance with RNAseq. We thank the Electron Microscopy Core Facility and Yinhui Lv for help in sample processing and TEM and SBF-SEM imaging. We also thank the Bioimaging AFM facility and Nigel Hodson for assistance with AFM. **Funding:** MRC project grants MR/T016744/1 and MR/P010709/1 (QJM, JAH); Versus Arthritis Senior Fellowship Award 20875 (QJM); Wellcome Trust Grant for the Wellcome Centre for Cell-Matrix Research 088785/Z/09 (QJM); BBSRC sLoLa grant (BB/T001984/1 to KEK and QJM); a RUBICON secondment fellowship to HM (European Union funded project H2020-MSCA-RISE-2015_690850); MRC DTP Studentship (HM). Research Grants Council of Hong Kong GRF17126118 (DC)

## Supplementary figures

**Figure S1.**
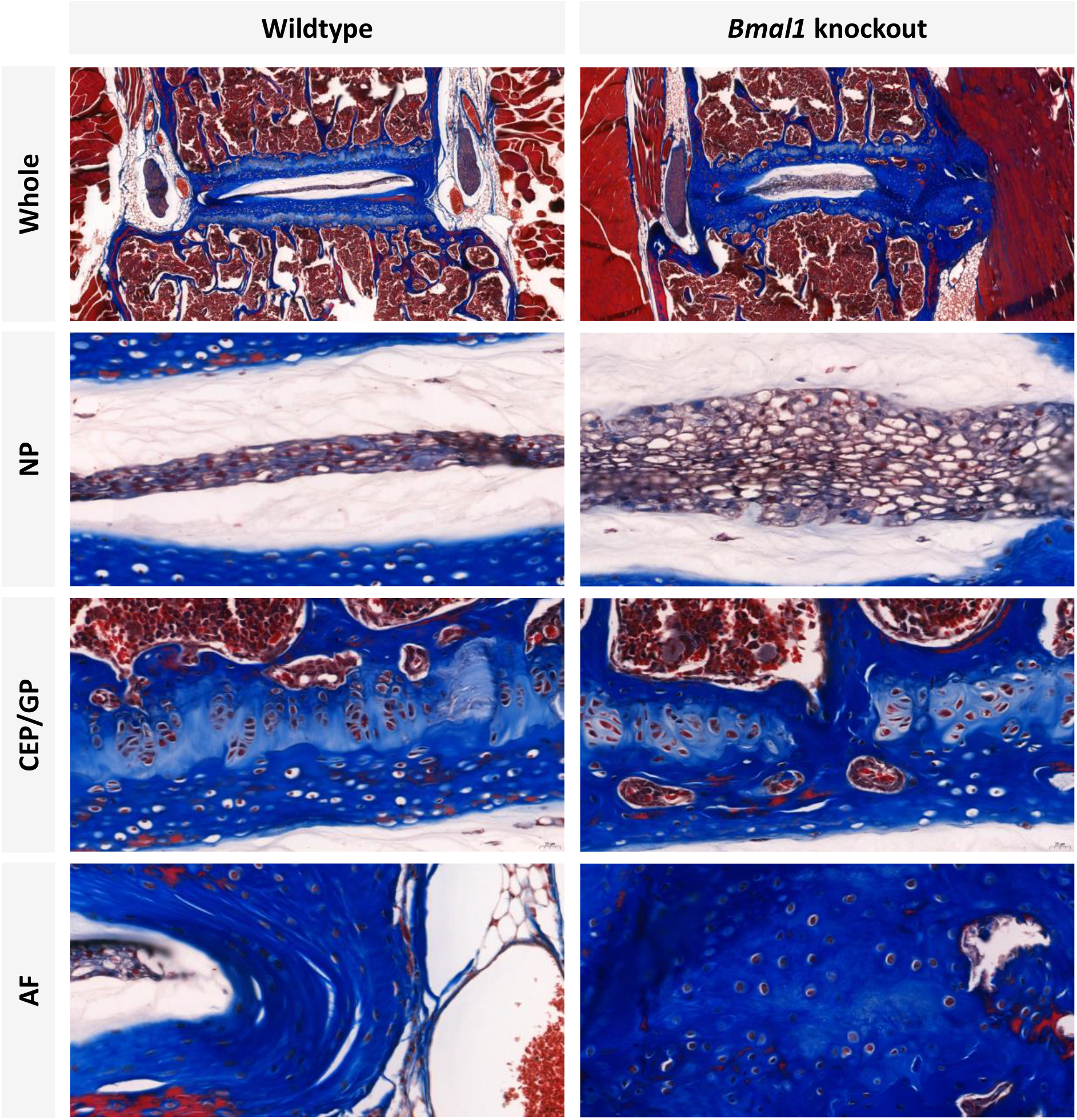
Masson’s trichrome staining of 12-month-old WT versus *Col2a1*-*Bmal1* KO lumbar intervertebral disc. *Bmal1* deletion resulted in a profound sclerotic phenotype in the aged disc, characterised by GP/CEP calcification and fusion of the vertebral bodies by bony tissue. N=4.

**Figure S2.**
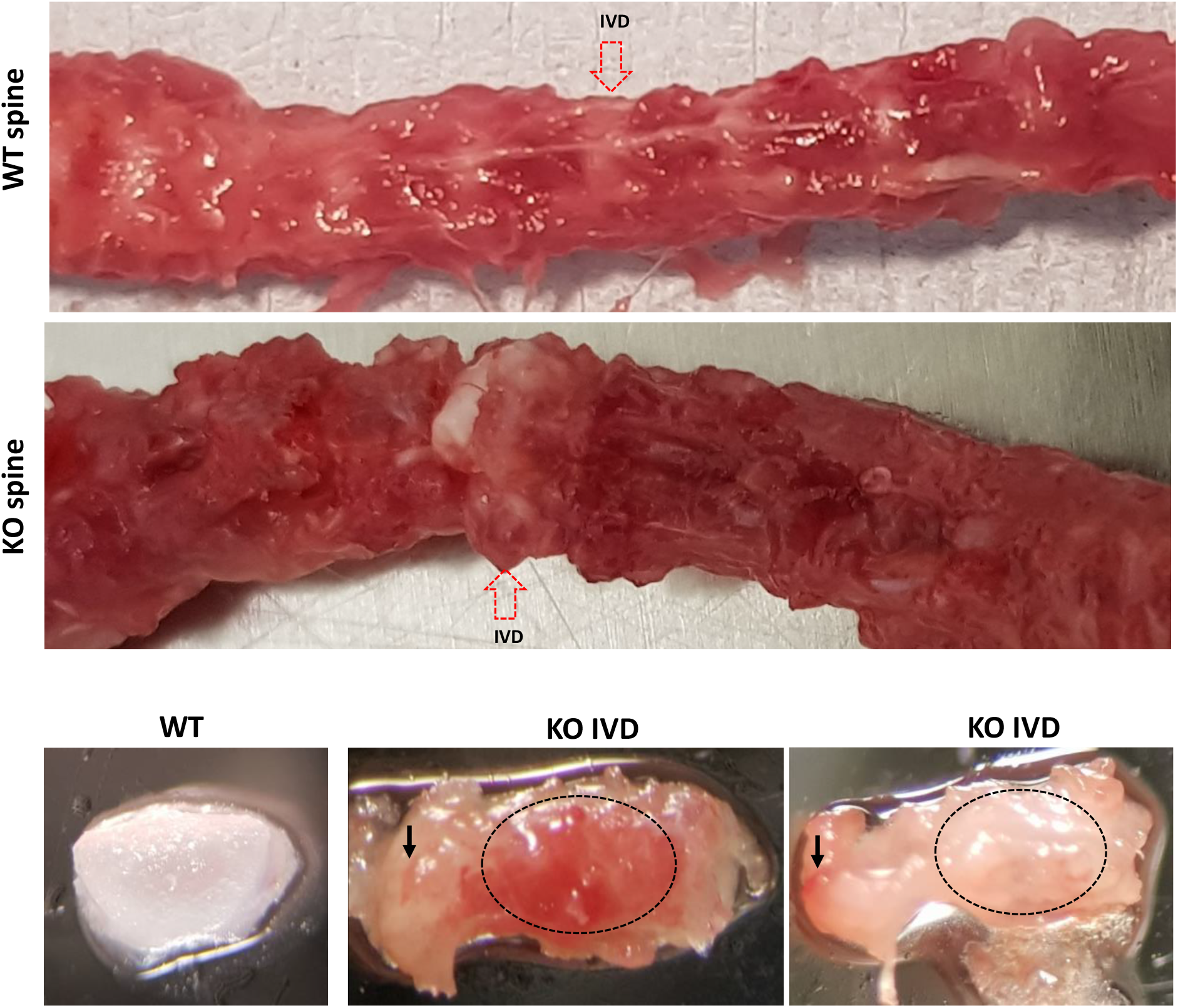
Dissected spines and macroscopic view of IVDs from 12-month-old WT and *Col2a1*-*Bmal1* KO. Arrows point to the corresponding IVDs in WT and *Bmal1* KO spine (top panels). Cartilaginous outgrowths in the outer AF are visible in resected IVDs of the *Bmal1* KO mouse. Dotted line demarks approximate location of the IVD. Black arrows point to possible ectopic calcification with red bone-like appearance. Cartilaginous end plate was not visible in *Bmal1* KO IVDs (bottom panels).

**Figure S3.**
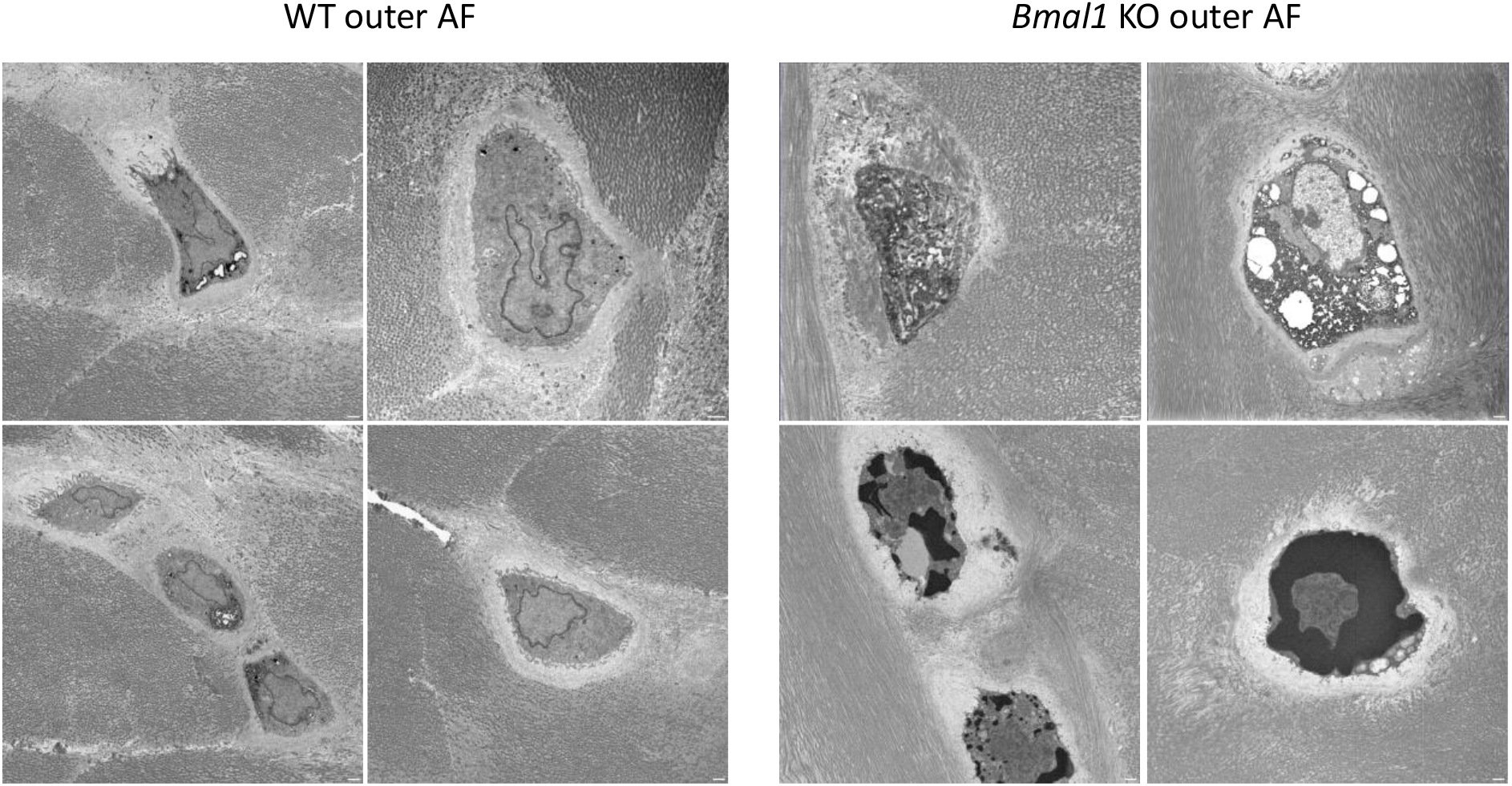
Examples of cell morphology in the outer AF of 12-month-old WT and *Col2a1*-*Bmal1* KO mice imaged by SBF-SEM. WT cells showed characteristic fibroblast like cell shape. Many *Bmal1* KO cells showed rounded chondrocyte morphology with electron dense regions in the cytoplasm. *Bmal1* KO cells were often surrounded by disorganised collagen fibrils. Scale bar=1µm.

**Figure S4.**
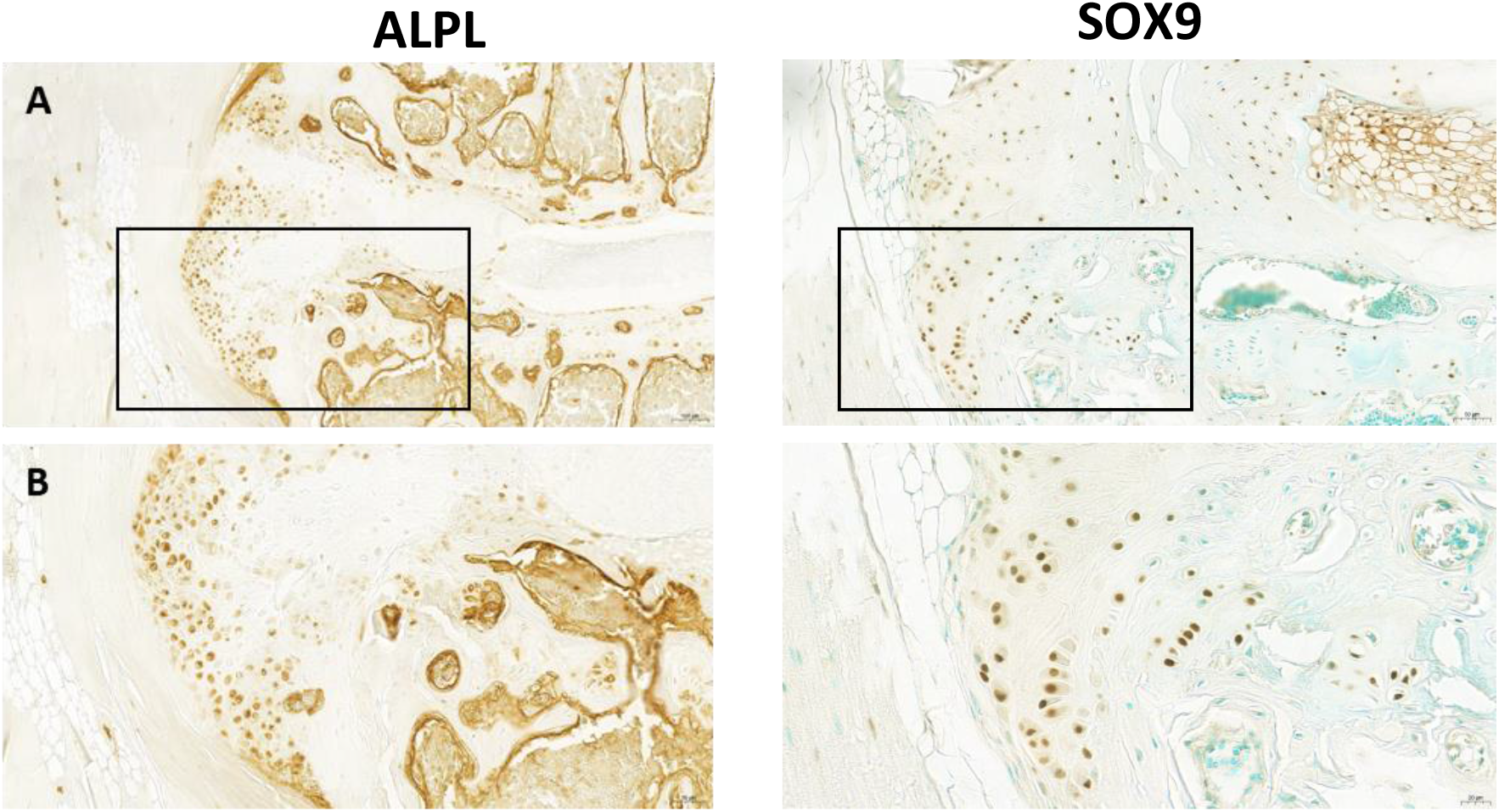
Immunohistochemistry staining of hypertrophic chondrocyte markers in the peripheral edge of the heterotopic ossification region in 12-month-old *Col2a1-Bmal1* KO lumbar intervertebral discs. **A)** ALPL and SOX9-positive cells were prolific throughout this region, particularly towards the peripheral edges. Cells showed a rounded morphology. **B)** x40 magnification regions highlighted in A).

**Figure S5.**
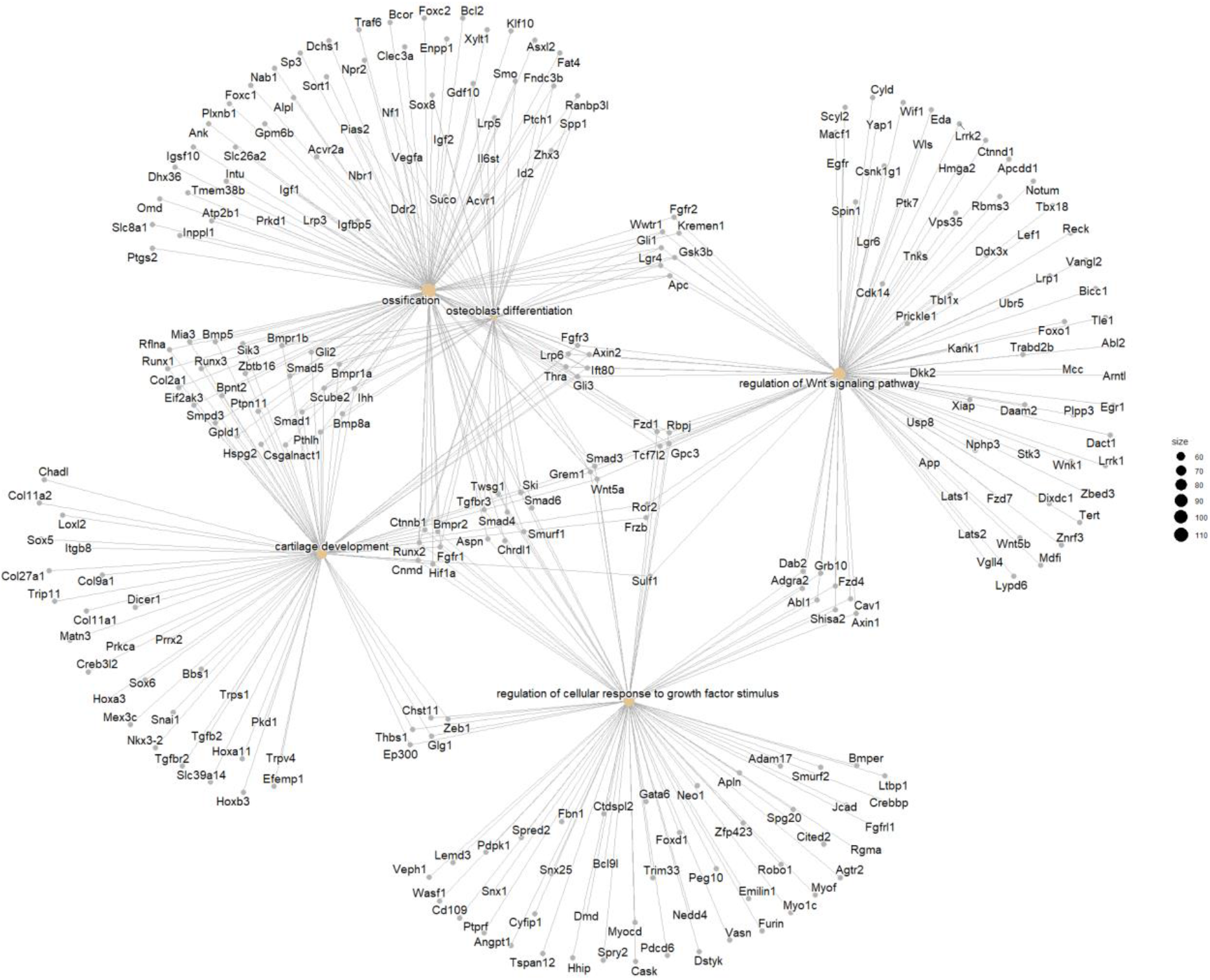
Detailed view of genes belonging to selected GO terms enriched in the 5 am/ 5 pm peaking ultradian transcriptome.

**Figure S6.**
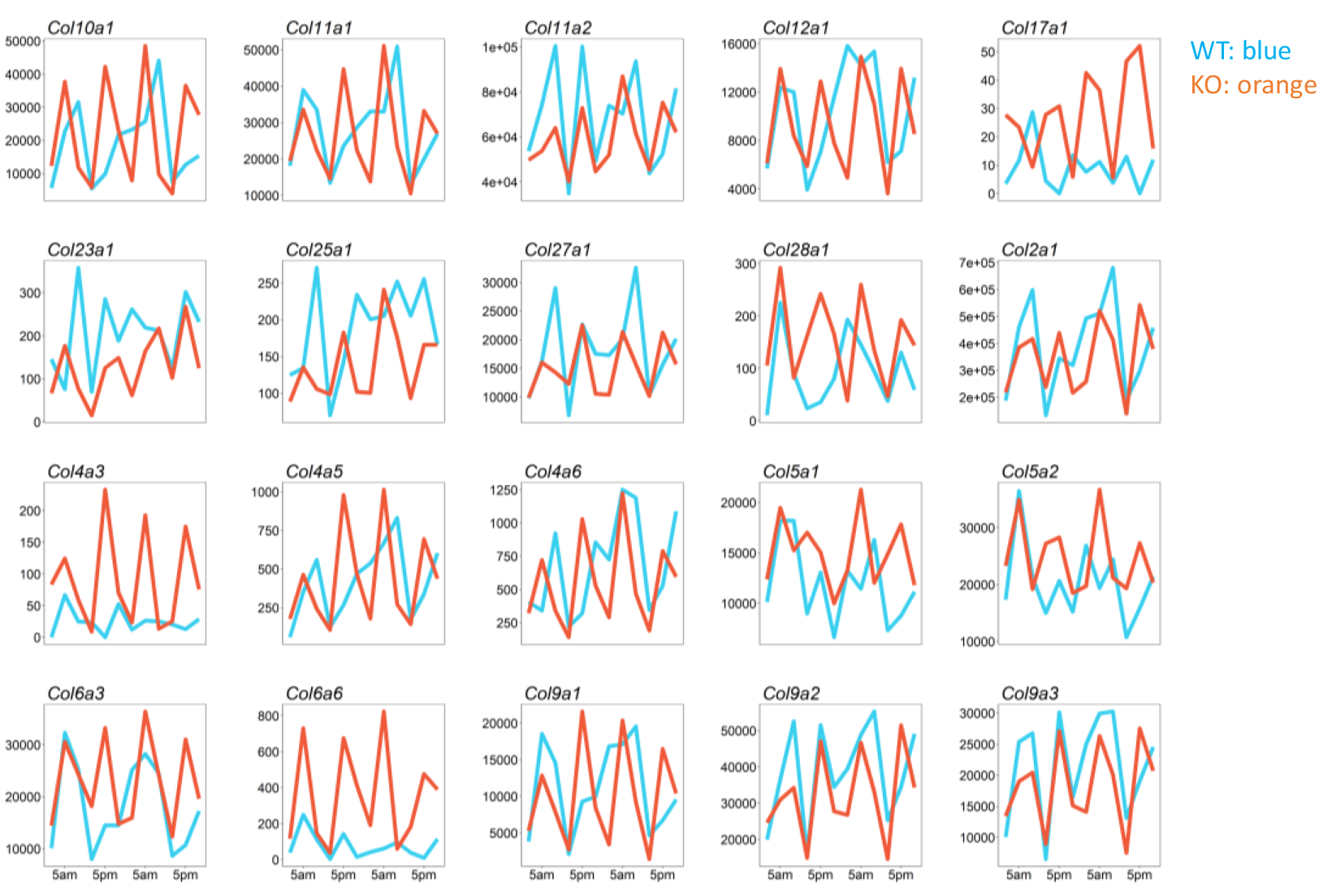
Ultradian collagen genes in the *Bmal1* KO IVDs and their corresponding expression in WT. Traces show wildtype (blue) and *Bmal1* KO (orange) gene expression over the 24-hour period studied. Y axis shows total read counts of each transcript.

**Figure S7.**
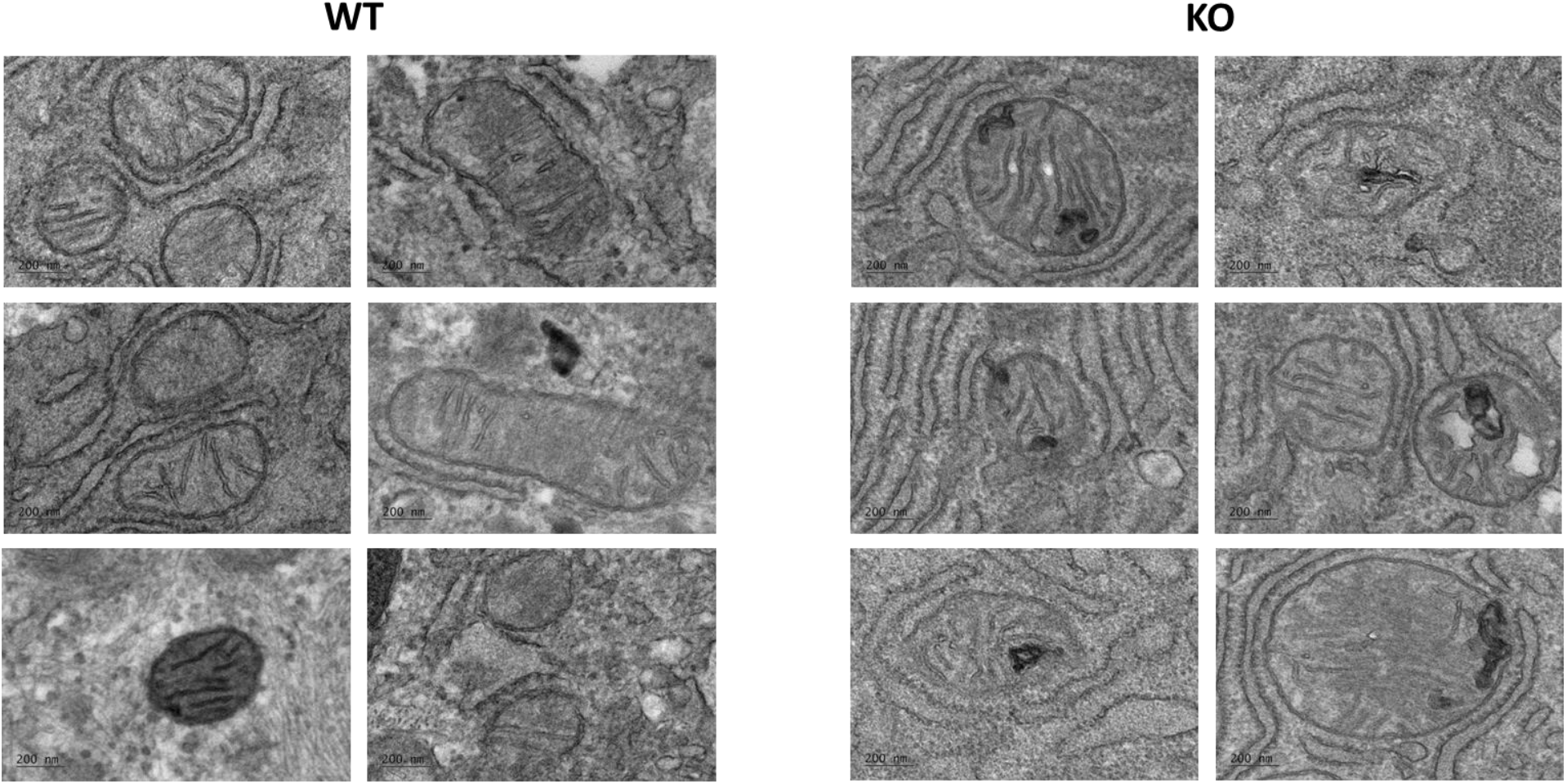
TEM images of mitochondria from the outer AF of 12-month-old wildtype and *Col2a1*-*Bmal1* KO mice. Electron-dense inclusions were often visible within the ***Bmal1*** KO mitochondria.

**Figure S8.**
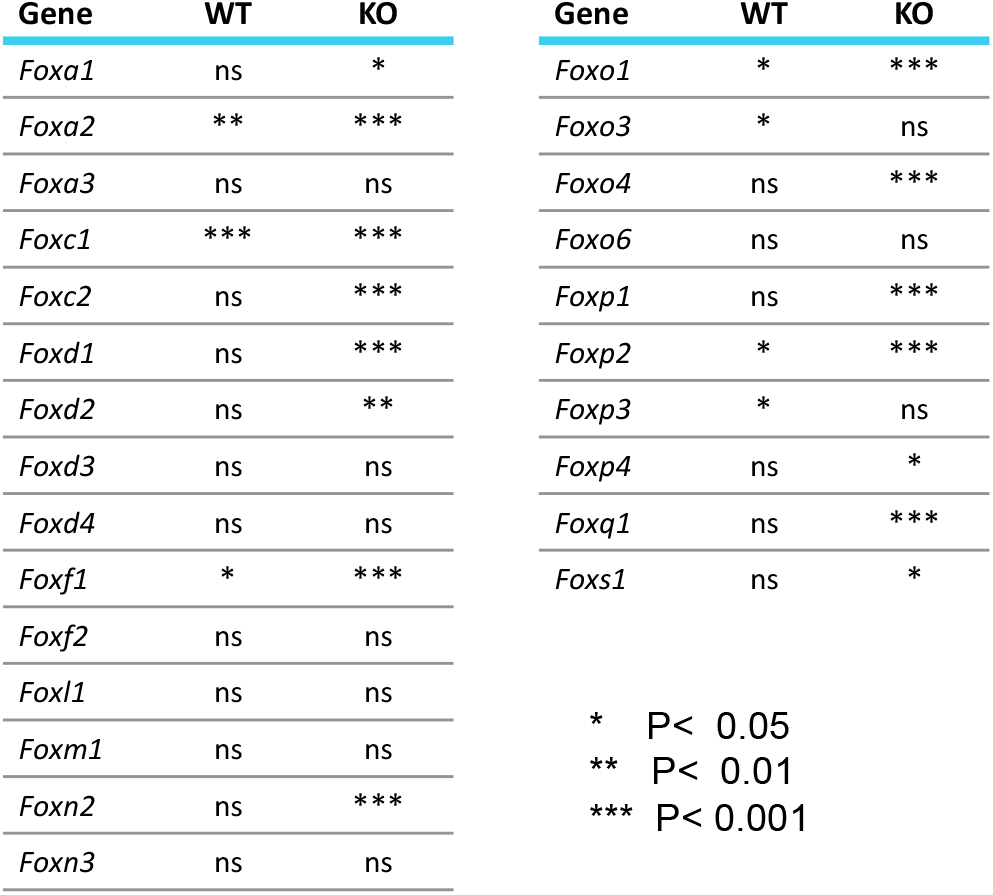
Ultradian expression of Fox family transcription factors. Fox factors are significantly more ultradian (12hr) in KO IVDs, as assessed by RAIN algorithm.

**Figure S9.**
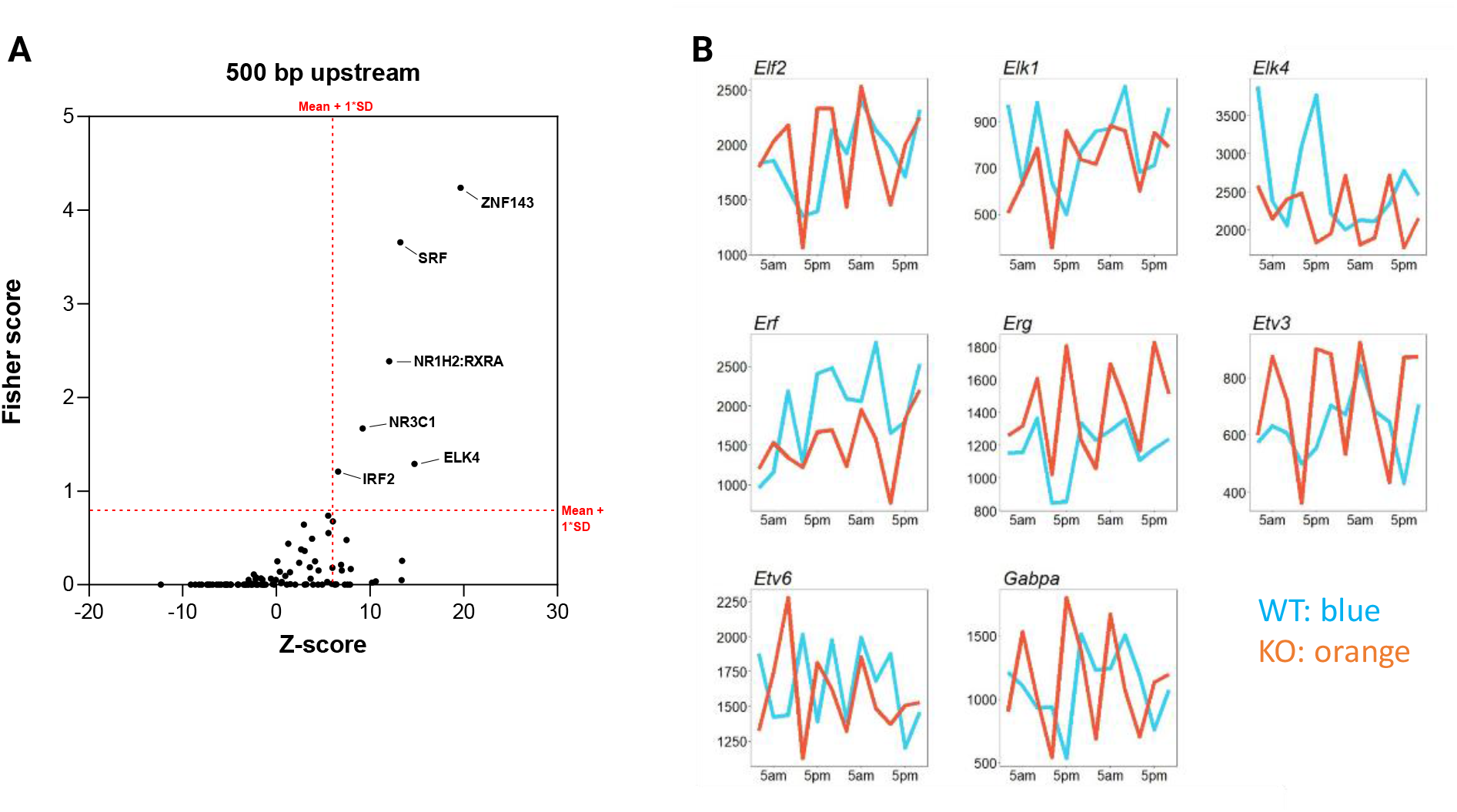
Upstream promoter analysis for the identification of over-represented transcription factor binding sites (TFBS) in the ultradian transcriptome of the *Bmal1* KO IVDs. **A)** Scatter plot of individual TFBS plotted against the Fisher scores and Z-score rankings of their enrichment in the regions 500 bp upstream of the transcription start sites of the 1 am/1 pm peaking ultradian transcriptome. **B)** Traces showing wildtype (blue) and *Bmal1* knockout (orange) gene expression of ETS transcription factor genes with significant ultradian rhythmicity in the *Bmal1* KO IVDs.

**Figure S10.**
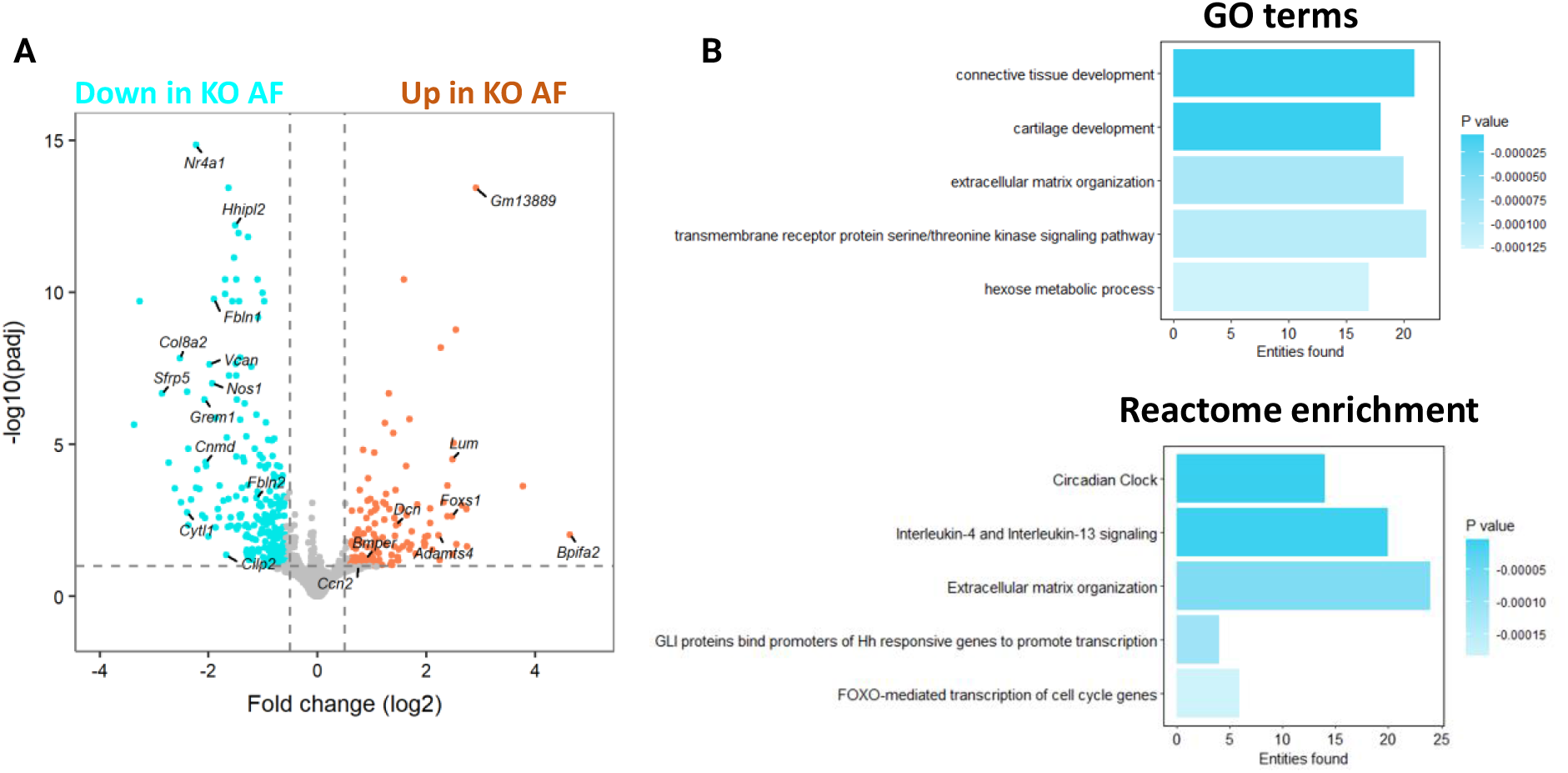
Differential gene expression in the AF of 3-month-old wildtype and *Col2a1*-*Bmal1* KO mice. **A)** Volcano plot showing differentially expressed genes in the AF between WT and *Bmal1* KO IVDs collected at 8 am. Log_2_FC 0.5 and adjusted p < 0.1 significance cut off. **B)** GO term (top) and Reactome pathway (bottom) enrichment analysis of WT versus *Bmal1* KO AF.

**Figure S11.**
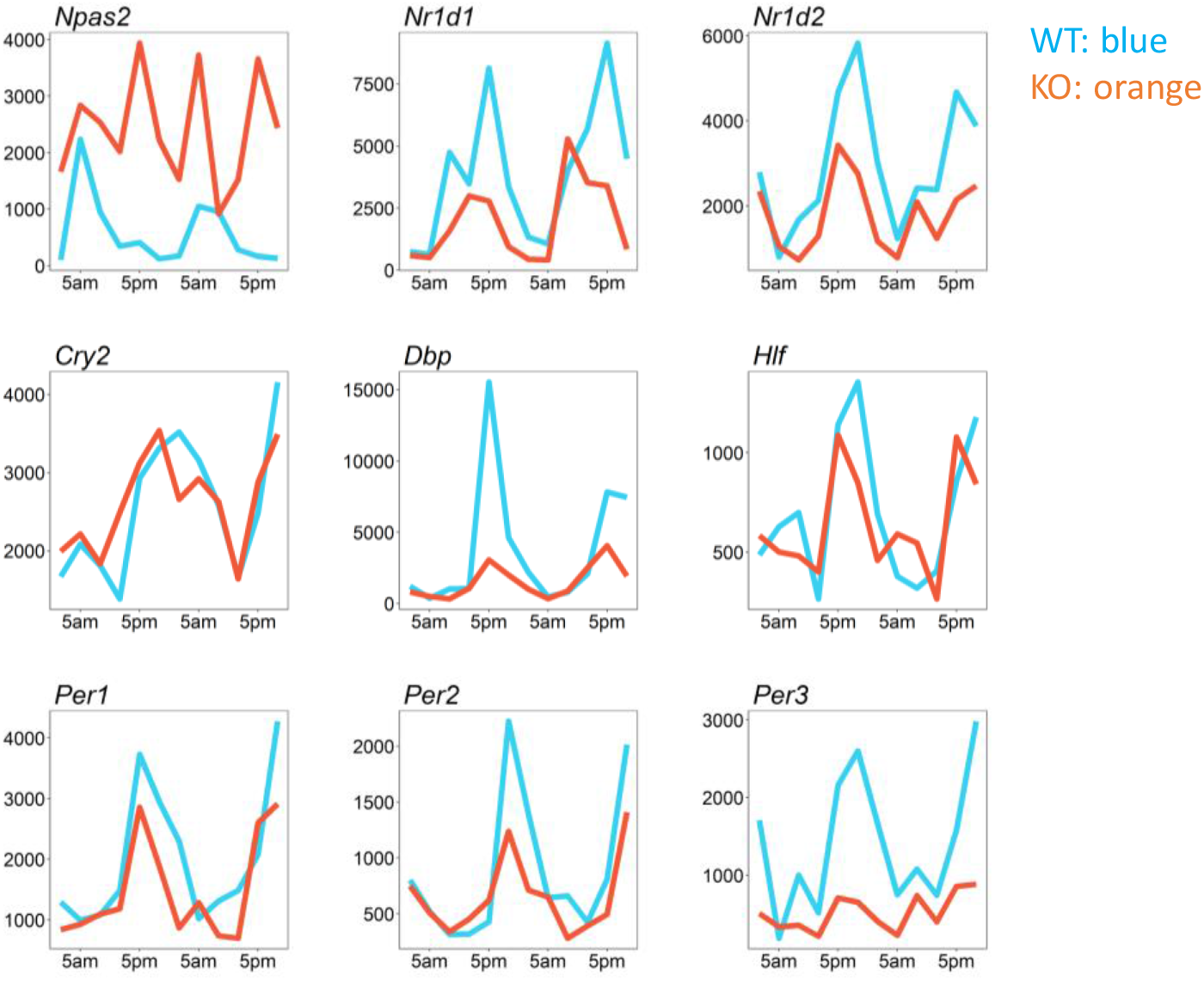
Gene expression profiles of core circadian clock genes in the wildtype and *Bmal1* KO IVDs. Traces show WT (blue) and *Bmal1* KO (orange) gene expression over the 24-hour period studied. Y axis shows total read counts of each transcript.

**Supplementary videos 1 and 2:** Electron micrograph of individual frames taken from 3view reconstructions of IVD tissues from WT (video 1) and *Bmal1* KO (video 2) mice.

### Supplementary Materials and Methods

#### Animals

All animal studies were performed in accordance with the 1986 UK Home Office Animal Procedures Act. Approval was provided by the local ethics committee. Mice were maintained at 20-22°C, with standard rodent chow available *ad libitum* and under 12:12 hr light dark schedule (light on at 7 am; light off at 7 pm). *Bmal1*^*flox/flox*^ mice^1^ were crossed onto a PER2::Luc background. The PER2::Luc mice carry the firefly luciferase gene fused in-frame with the 3’ end of the *Per2* gene, creating a fusion protein reporter^2^. *Bmal1*^*flox/flox*^ - PER2::Luc mice were subsequently crossed with *Col2a1-Cre* mice expressing Cre recombinase under the control of the *Col2a1* promoter^3^ to generate IVD /cartilage specific *Bmal1* KO. All mice were bred in-house at the University of Manchester. Generation and genotyping of the *Col2a1-Bmal1*^*-/-*^ mice was described before^4^.

#### Histology and immunostaining

Lumbar spines from 12-month-old mice were fixed in 4% PFA for 24 hours, followed by decalcification in 10% EDTA pH 7.4 for 3 weeks at room temperature. Decalcified tissues were subsequently dehydrated, embedded in paraffin, and sectioned at a thickness of 5 µm. For histological staining, Masson’s trichrome staining (Abcam) and RGB trichrome staining^5^, tissue sections were sequentially deparaffinised, rehydrated and stained according to the manufacturer’s and authors’ methodologies.

For immunohistochemistry, tissue sections were sequentially deparaffinised, rehydrated and equilibrated in PBS. Endogenous peroxidase was quenched by incubating slides in PBS with 1% hydrogen peroxide (Sigma) for 10 minutes. Antigen retrieval was then performed using 1 mg/ml trypsin (Sigma) in PBS for 10 minutes at room temperature. Sections were washed and non-specific binding was blocked in blocking solution (3% serum in PBS with 0.1% Triton X) for 1 hour at room temperature. Primary antibodies for ALPL (R&D Systems, AF2910), SOX9 (Cell Signalling) were diluted at 1:1000 and 1:500, respectively, in blocking buffer. Sections were incubated in this solution overnight at 4°C in a humidified chamber. Slides were then washed thoroughly and incubated in biotinylated secondary antibody (Vector Labs, BA-1000/BA-9500) diluted 1:2000 in blocking solution for 1 hour at room temperature, followed by incubation in HRP Streptavidin (Vector Labs, SA5004) for 1 hour at room temperature. Peroxidase signal was developed with DAB substrate (Merck Millipore) for 2–3 minutes. SOX9-stained sections were lightly counterstained using methyl green (Sigma). Sections were washed, sequentially dehydrated and mounted in DPX mounting media. Stained slides were scanned by a Pannoramic 250 slide scanner (3DHistech) and representative images extracted using CaseViewer v2.4 software (3DHistech).

#### Electron microscopy (EM)

AF tissues were resected under dissection microscope from lumbar IVDs from 12-month-old WT and *Col2a1-Bmal1*^*-/-*^ mice. Tissues were then fixed in 1% osmium and 1.5% potassium ferrocyanide in 0.2 M sodium cacodylate buffer for 1 hour, washed with distilled water, incubated with 1% tannic acid in 0.1 M cacodylate buffer for 1 hour, washed with distilled water and then incubated with 1% osmium tetroxide in water for 30 minutes. The samples were then washed with distilled water and stained with 1% uranyl acetate in water for 1 hour, dehydrated in acetone and embedded in resin. TEM and SBF-SEM imaging was performed as described previously^6^.

#### Atomic Force Microscopy (AFM)

Unfixed and non-decalcified lumbar IVDs from 6-week-old and 12-month-old mice were cryosectioned at a thickness of 20 µm using a Feather C35 Carbon Fibre Blade (PFM Medical) on a CM3050S Cryostat (Leica). Sections were mounted onto SuperFrost Plus slides (Thermo Fisher Scientific). Prior to AFM imaging, sections were washed three times in ultrapure water and left overnight to dry fully. A BioScope Catalyst AFM (Bruker) was mounted onto an Eclipse Ti-U inverted microscope (Nikon) and fitted with ScanAsyst Air cantilever (Bruker). Imaging was performed using NanoScope v9.1 software (Bruker) in ScanAsyst mode in air at a scan rate of 0.399 Hz and 512 samples/line. Peak force error images were used to quantify fibril diameter and length of D periods, using NanoScope Analysis v1.5 software (Bruker).

#### Tissue collection and RNA-sequencing

For circadian time series whole lumbar IVDs from 2-3-month-old male and female WT and *Col2a1-Bmal1*^*-/-*^ mice were collected 4 hours apart over a period of 48 hours, starting at 9 am. Three biological replicates were obtained per genotype and timepoint. Whole lumbar IVDs were resected, and paraspinal tissues carefully removed. Tissues were immediately snap frozen and stored at -80°C for later processing. For AF tissue RNAseq male and female three-month-old WT and *Col2a1-Bmal1*^*-/-*^ mice were collected at 8 am. Three biological replicates were obtained per genotype and timepoint. Tail discs were resected, and the NP and AF were manually separated under a dissection microscope to ensure careful separation of tissue. AF tissues were kept on ice and kept hydrated in HBSS throughout the dissection procedure, then snap frozen and stored at -80°C for later processing.

For the isolation of RNA tissues were homogenised in TRIzol using a Mikro-Dismembrator S (Satorius Stedim Biotech) at 2000 rpm for 30 seconds, twice. Liquid nitrogen was used to keep tissues and homogenates cold at all times during the homogenisation process. Homogenates were subsequently processed using 1-Bromo-3-chloropropane to separate RNA. The aqueous phase was processed twice to ensure clean separation of RNA from other organic matter. Glycogen was used to capture mRNA. Pellets were washed three times in 70% EtOH, dried and resuspended in RNAse-free H_2_O. The quantity of resuspended mRNA was measured using a Qubit™ 4 Fluorometer (Thermo Fisher Scientific) and RNA integrity measured using a 2200 TapeStation (Agilent Technologies) to ensure good quality for downstream analysis.

Libraries were generated by the Genomic Technologies Core Facility using the TruSeq® Stranded mRNA assay (Illumina, Inc.) according to the manufacturer’s protocol. Briefly, total RNA (0.1-4 μg) was used as input material from which polyadenylated mRNA was purified using poly-T, oligo-attached, magnetic beads. The mRNA was then fragmented using divalent cations under elevated temperature and then reverse transcribed into first strand cDNA using random primers. Second strand cDNA was then synthesised using DNA Polymerase I and RNase H. Following a single ‘A’ base addition, adapters were ligated to the cDNA fragments, and the products then purified and enriched by PCR to create the final cDNA library. Adapter indices were used to multiplex libraries, which were pooled prior to cluster generation using a cBot instrument. The loaded flow-cell was then paired-end sequenced (76 + 76 cycles, plus indices) on an Illumina HiSeq4000 instrument. Finally, the output data was demultiplexed (allowing one mismatch) and BCL-to-Fastq conversion performed using Illumina’s bcl2fastq software, version 2.20.0.422.

Unmapped paired-end sequences were tested by FastQC (http://www.bioinformatics.babraham.ac.uk/projects/fastqc/). Sequence adapters were removed and reads were quality trimmed using Trimmomatic_0.39^7^. The reads were mapped against the reference mouse genome (mm10/GRCm38) and counts per gene were calculated using annotation from GENCODE M21 (http://www.gencodegenes.org/) using STAR_2.7.7a^8^. Normalisation and differential expression was calculated with DESeq2_1.30.1^9^.

#### Analysis of temporal RNAseq data

Circadian time series RNA-seq data was analysed using the RAIN algorithm, a non-parametric method for detecting waveform patterns in biological datasets^10^. Waveforms showing a period of 24 or 12 hours were identified and those which met the selection threshold (P < 0.01) were selected for further downstream analysis. Rhythmic genes were clustered by non-hierarchical clustering using the pheatmap package for R and plotted as heatmaps arranged by waveform shape similarity. For AF RNAseq analysis genes were labelled as being differentially expressed if they passed thresholds with both an FDR-corrected p-value (padj) of 0.1 and a Log2FC of 0.58 (i.e., a 1.5-fold change).

GO term enrichment was performed with the ClusterProfiler package for R. Enriched GO terms with a significance threshold of P < 0.01 were selected. The ‘simplify’ function was then applied to remove redundancy in GO terms based upon their semantic similarity. Reactome pathway analysis was performed using the Reactome Knowledgebase (https://reactome.org).

Upstream regulator analysis was performed using Opossum v3. The sequences of genes identified as being significantly rhythmic were analysed against a background of the non-rhythmic intervertebral disc transcriptome for enriched transcription factor binding site sequences 500 bp upstream of the transcription start site. Transcription factor binding site sequences were to be considered significantly enriched when they showed both a Z-score and Fisher score above the mean +1*SD score for all transcription factors within the JASPAR database.

#### Gene expression analysis

Total RNA was isolated and purified using the PureLink RNA Mini Kit (Ambion) according to manufacturer’s instructions. Purified RNA was quantified using a NanoDrop 2000 spectrophotometer (Thermo Fisher) and transcribed to cDNA with the High-Capacity RNA-to-cDNA kit (Applied BioSystems). TaqMan-based qPCR was performed using a StepOnePlus Real-Time PCR System (Applied Biosystems, Thermo Fisher Scientific) with FAST Blue qPCR MasterMix (Eurogentec) and TaqMan primer-probe single-tube assays (Applied Biosystems): *Foxo1* (Mm00490671_m1)

#### Osteogenic differentiation

Human AF-S cell line^11^ were differentiated in osteogenic induction media containing 10% FBS (Gibco), 50 μg/ml 2-phospo-L-ascorbic acid trisodium salt (Sigma-Aldrich) and 100 nM Dexamethasone (Sigma-Aldrich) in DMEM/F-12 (Gibco). Media were replenished every 2-3 days for a duration of 2 to 3 weeks. For differentiation experiments involving FOXO1 inhibitor AS1842856 (Calbiochem), the compound was present in media for the full duration of osteogenic induction at a dose of 0.5 μM.

